# Body size dependent dispersal influences stability in heterogeneous metacommunities

**DOI:** 10.1101/2021.02.08.430322

**Authors:** Kurt E. Anderson, Ashkaan K. Fahimipour

**Affiliations:** University of California Riverside, Department of Evolution, Ecology, & Organismal Biology; National Oceanic and Atmospheric Administration, Southwest Fisheries Science Center; University of California Davis, Department of Computer Science

## Abstract

Body size affects key biological processes across the tree of life, with particular importance for food web dynamics and stability. Traits influencing movement capabilities depend strongly on body size, yet the effects of allometrically-structured dispersal on food web stability are less well understood than other demographic processes. Here we study the stability properties of spatially-arranged model food webs in which larger bodied species occupy higher trophic positions, while species’ body sizes also determine the rates at which they traverse spatial networks of heterogeneous habitat patches. Our analysis shows an apparent stabilizing effect of positive dispersal rate scaling with body size compared to negative scaling relationships or uniform dispersal. However, as the global coupling strength among patches increases, the benefits of positive body size-dispersal scaling disappear. A permutational analysis shows that breaking allometric dispersal hierarchies while preserving dispersal rate distributions rarely alters qualitative aspects of metacommunity stability. Taken together, these results suggest that the oft-predicted stabilizing effects of large mobile predators may, for some dimensions of ecological stability, be attributed to increased patch coupling *per se*, and not necessarily coupling by top trophic levels in particular.

## Introduction

What allows large, complex ecosystems to be stable? May’s analysis of randomly arranged communities of self-limiting populations challenged previous ecological thinking on this issue, showing that greater species richness and interaction connectivity tended to destabilize random communities rather than stabilize them [1, 2]. This and subsequent theory set the stage for decades of work analyzing the complicated relationship between diversity and dynamics that continues today [3, 4, 5, 6, 7]. In contrast to randomly-assembled communities, an emerging focus in modern biodiversity theory is on the non-random structural features of ecological systems that impart stability [4, 8, 9]. For communities organized around feeding relationships (*i.e.,* food webs), two types of *structure* receiving extensive attention are allometric hierarchies, where larger species mostly eat smaller ones and populations experience other demographic rates dependent on body size [5, 10, 11]; and dispersal among spatially discrete habitats [12, 13, 14, 15].

Body size-based food web topologies and allometric scaling of population demographic rates have both shown to be stabilizing for models of trophic interactions [5, 6, 10, 16, 17]. Simple mass-based hierarchical feeding rules — where species high in the feeding hierarchy are interpreted as large-bodied predators — can successfully reproduce realistic food web topologies [4, 5, 11, 18, 19, 20, 21] that are more likely to be dynamically stable than random network configurations [16, 17]. Likewise, allometric scaling of species’ demographic rates such as handling times, conversion efficiencies, and biomass turnover, are predicted to stabilize food webs [5, 10, 22, 23, 24, 25]. This is particularly the case when the ratio of body masses between resources and consumers is large and consistent with values observed in natural food webs [5, 10, 26].

At larger scales, dispersal generates structure by linking spatially distinct food webs through the movement of individuals. The role of dispersal in population dynamics and community composition is a central focus in ecology, with early work emphasizing the colonization of islands by mainland species [27, 28] and the “rescue” of small populations in sink habitats [29]. Colonization-extinction dynamics in spatially subdivided habitats, originally examined in a population context [30, 31], extended these results and have been shown to promote higher regional food web diversity than can be supported in isolated well-mixed systems [15]. Dispersal can also stabilize species interactions locally by mimicking density-dependence in *per capita* growth rates [32, 33, 34, 35]. More recent work has linked metacommunity dynamics to May’s original examination of species richness and connectance, showing that dispersal among spatially distinct subpopulations can be strongly stabilizing [36, 37]. Complexity-stability relationships in these cases are relaxed or even reversed relative to results seen in linear stability analyses of random matrices.

Much like trophic interaction rates, many traits that influence animal locomotion, movement speed, and potentially dispersal also vary with body size [38, 39, 40]. Spatial patterns of resource use, home ranges, and geographic range size also exhibit strong allometric relationships [24, 41, 42, 43, 44]. These patterns in turn create the potential for spatial coupling of habitat patches [45] and hence local food webs [16, 46, 47] that depends on body size. While many studies emphasize faster movements of large consumers, suggesting positive body size dispersal relationships, large-bodied species many face greater dispersal limitation in some habitats [48, 49, 50], suggesting a range of potential relationships between body size and dispersal across ecosystems [15].

Theoretical and empirical evidence suggests an important role for body size-dispersal scaling in ecosystem dynamics, yet the full effects of dispersal variation among species in food webs have so far been difficult to systematize [15]. Some mathematical models indicate that the coupling of distinct food webs by consumer movement can be stabilizing when those webs represent different energy channels or environmental conditions [12, 16, 47]. In other models, greater mobility of consumers is a key requirement for instability [51, 52, 53, 54]. Yet other examples have identified dispersal-driven instabilities for communities in which primary producers traverse space more rapidly than other species [13]. Overall, the effects of dispersal are complicated in ecological networks and a general understanding of how dispersal rules influence food web stability is lacking [13, 15].

Here, we examine how body size scaling of species’ dispersal rates influences stability in model trophic metacommunities (Fig. 1). In particular, we ask whether body size-dependent variation in dispersal rates influences trophic metacommunity stability relative to rates that are either uniform or randomly varying among species. We examine landscapes of discrete habitat patches that include variation in local abiotic conditions, generating spatial heterogeneity in rates of primary production and trophic interactions among species. Clearly, the stability of such heterogeneous metacommunities will depend on the proportion of patches in the landscape with locally favorable conditions for stability. We show how the body size scaling of species’ dispersal rates alters this relationship. Because general rules describing the dependence of dispersal on body size are lacking and likely vary among ecosystems [15], we consider both positive and negative relationships between dispersal rates and body sizes. Our results show strong effects of dispersal-body size scaling on metacommunity stability, largely due to increased connectivity among local webs with different stability properties.

**Figure 1:**
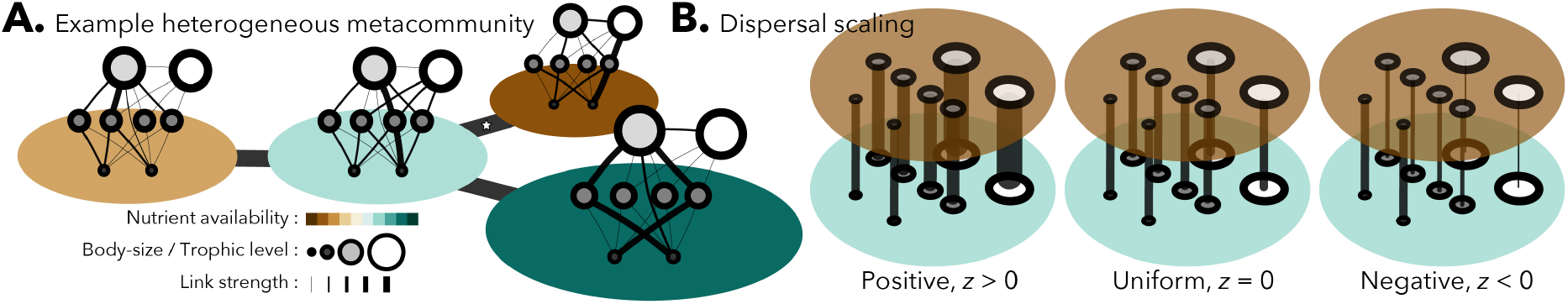
Model metacommunities are composed of local food webs connected to one another by dispersal. **A.** Each local web inhabits a habitat patch that is part of a spatial network, generated as a random geometric graph. Species in each web have a body size that is larger at higher trophic levels. Food webs have the same number of species and topology in all patches, but interaction rates and other ecological parameters vary among habitats mimicking spatial environmental heterogeneity. **B.** Dispersal varies as either an increasing or decreasing function of body size.

## Model formulation

We model trophic metacommunities as copies of food webs consisting of *S* species, embedded in a set of *N* patches. We chose to represent spatial networks as random geometric graphs, which provide a reasonable approximation for real networks [15] of habitats and the dispersal connections between them (Fig. 1).

Food web topologies were generated using the niche model [18], which recapitulates realistic yet variable feeding relationships with only two input parameters, species richness *S* and connectance *C* [18]. Briefly, species are assigned a position on a 1-dimensional niche axis and feed on species over a range determined by *C*. The feeding range is centered below the species’ niche positions, creating a trophic hierarchy where each species *i* has trophic position *T_i_*, defined by the path length to any basal producer.

Following [5, 7, 10, 11], we assigned body sizes assuming that the normalized mass *M_i_* of each species *i* scales with their trophic position *T_i_*, as *M_i_* = *R^T_i_^*. For reported results, normalized producer body sizes are uniformly set to one while *R* = 42. This value of R represents the average predator-prey body mass ratio reported by [26], although our qualitative results are consistent across a wide range of realistic ratios.

Dynamics on the links defined by each niche model topology were represented as the set of ordinary differential equations of the form

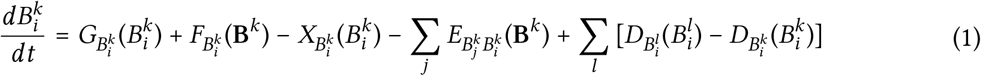

where 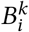 is the biomass density of species *i* in patch *k.* The non-specified function *G* is the growth rate of primary producers, *F* is the rate of biomass accumulation due to feeding on other species, *X* is rate of biomass loss due to respiration and mortality, *E* is the rate of biomass loss due to consumption by species *j,* and *D* is the dispersal rate between patches. While the functional forms are not explicitly specified, the general form of equation (1) admits the calculation of a Jacobian matrix that quantifies how species in the metacommunity respond to perturbations from steady state and therefore metacommunity stability [55, 56]. Using the generalized modeling method [6, 55, 57], the derivatives of functions *G, F, X*, and *E* that constitute the non-dispersal elements of the Jacobian matrix can be recast in terms of scale, branching, and elasticity parameters (see Methods). The generalized model parameters have clear ecological interpretations: scale parameters set the time scale of biomass turnover, while branching and elasticity parameters set the relative contributions of different processes to biomass gains and losses and the form of non-linearities, respectively [6, 55, 56, 57]. The range of generalized model parameter values studied here map to most commonly encountered functional forms (*e.g.,* Lotka-Volterra-like and Holling type II and III functional responses) that link interaction rates with species’ densities in conventional models, and follow ecologically-based arguments from prior work [6, 55, 56] (Table 1).

**Table 1.** Parameter definitions and ranges used in metacommunity simulations.

| Parameter | Description | Range or Value |
| --- | --- | --- |
| **Scale** |  |  |
| $\alpha_i$ | Normalized turnover rate of species $i$ | $M_i^{-1/4}$ |
| **Branching** |  |  |
| $\rho_i^k$ | Fraction of biomass gains from predation, for species $i$ in patch $k$ | 0 or 1 |
| $\sigma_i^k$ | Fraction of biomass loss from predation, for species $i$ in patch $k$ | [0, 1] |
| $\beta_{ij}^k$ | Fraction of species $i$'s loss from consumption by $j$ in patch $k$ | [0, 1] |
| $\chi_{ij}^k$ | Fraction of species $j$'s gains from consumption of $i$ in patch $k$ | [0, 1] |
| **Elasticity** |  |  |
| $\phi_i^k$ | Nutrient availability for producer $i$ in patch $k$ | [0, 1] |
| $\psi_i^k$ | Sensitivity of $i$'s predation rate to its own density in patch $k$ | [0.5, 1] |
| $\lambda_{ji}^k$ | Sensitivity of $j$'s foraging preferences to prey densities in patch $k$ | [1] |
| $\gamma_i^k$ | Sensitivity of $i$'s predation rate to total prey density in patch $k$ | [0.5, 1.5] |
| $\mu_i^k$ | Sensitivity of species $i$'s mortality to its own density in patch $k$ | [1, 1.5] |
| **Niche model** |  |  |
| $S$ | Species richness | [10, 30] |
| $C$ | Food web connectance | [0.12, 0.24] |

Trophic metacommunities were constructed as follows. First, a spatial structure of *N* = 10 habitat patches was randomly generated as a geometric graph on the interval [0,1] with a neighborhood radius of 0.32 (see Methods). Each patch *k* in the spatial network contains a copy of the same *S* species niche model [18] web. Heterogeneity in factors that influence primary production and trophic interactions were modeled as random variation among patches in branching and elasticity parameters [6, 55, 56, 57]. Values of these spatially variable parameters were drawn independently from uniform distributions defined by ecologically meaningful ranges (Table 1).

For simplicity, we assume that there is no dependence of *per capita* dispersal rates on patch identity or on interspecific densities (i.e. no cross-diffusion). We further assume that dispersal is a linear function of local intraspecific density and allow for dispersal rates to vary among species,

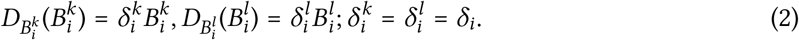

Each species *i* in a particular metacommunity scenario is assigned a species-specific dispersal rate *δ_i_* that is dependent on its body size. We chose a power law relationship between body mass and dispersal rate, as this general form captures many allometric scaling relationships related to locomotive capabilities and spatial habitat use [24, 38, 40, 58]. Specifically,

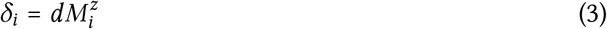

where *d* is the global link strength of the spatial network and *z* is the body size scaling exponent for dispersal. Producers have a body size equal to one such that for producers, *δ_i_* = *d*. When *z* is positive, species with higher trophic positions and thus larger body sizes traverse the spatial network at a higher rate. This scenario potentially reflects terrestrial and pelagic food webs where larger animals have greater mobility and hence dispersal potential [12]. In other systems, *z* may be negative [13, 15, 40], giving species in lower trophic levels the fastest dispersal rates with lower rates for larger bodied species.

We examined metacommunity stability using linear stability analysis. This procedure was applied in classic work on community stability by May and others [1, 2] and has since been expanded to incorporate spatial processes [14, 36]. Linear stability is assessed by examining the eigenvalues of the Jacobian matrix of the trophic metacommunity, **J**. Each local food web has a corresponding local Jacobian J_*k*_ that is derived from equation (1); these local Jacobians are collected and numerically arranged as blocks on the diagonal of the *SN* × *SN* matrix **P**. The local food web information is then used to calculate the metacommunity Jacobian **J** using the equation [13, 14, 59]

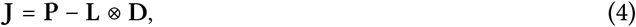

where ⊗ is the Kronecker product; the *N* × *N* matrix **L** is the Laplacian of the corresponding spatial network; and **D** is an *S* × *S* Jacobian-like diagonal matrix containing species-specific dispersal rates calculated from equation (2). Eigenvalues of local food web Jacobians **J**_*k*_ and the metacommunity Jacobian **J** were computed numerically; stability occurs when the real parts of all eigenvalues are negative. Additional details regarding this method of analysis are found in refs. [13, 14, 59, 60] and in the Methods.

Body size scaling could increase stability in metacommunities owing to simple increases in overall dispersal, rather than because of particular relationships between body size, trophic position and dispersal rate. To test this possibility, we compared our suite of model metacommunities with different dispersal rules to those with random variation in dispersal rates. For each metacommunity and each dispersal scenario, we computed Jacobians with 100 random permutations of species’ dispersal rates (*i.e.*, the diagonal entries of **D**, see eq. 3) and compared stability properties in these randomized metacommunities to their corresponding intact systems.

## Results

Linear stability of both local food webs and metacommunities varied dramatically over the multitude of food webs examined, depending on species richness, food web connectance, branching, elasticity [6], and dispersal parameters. The baseline dispersal rate *d* and body size scaling exponent *z* in particular influenced metacommunity stability. Figure 2 presents the aggregated results of all numerical analyses organized by local food web and metacommunity stability. The proportion of stable local food webs gives the proportion of webs in a metacommunity that would be stable in the absence of any dispersal. The proportion of stable metacommunities gives the probability that a corresponding trophic metacommunity will be stable when those patches are then linked by dispersal. The proportion of stable local food webs has the intuitive effect of positively increasing the chance that the metacommunity they constitute will also be stable; a metacommunity composed entirely of locally stable food webs is always stable. However, the strength of this positive relationship relies critically on the rules that determine dispersal rates across trophic levels.

**Figure 2:**
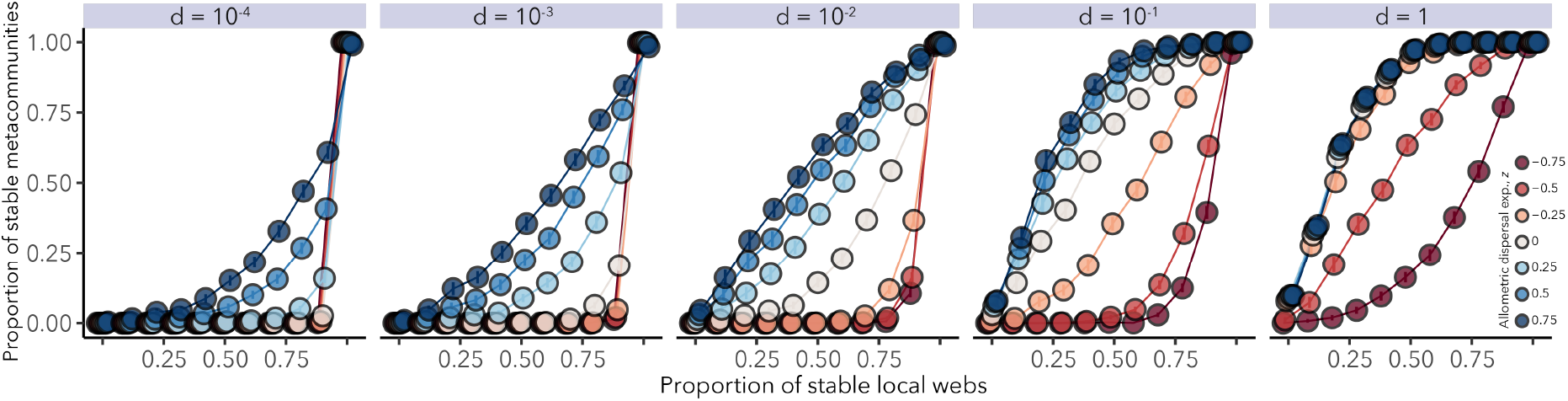
The relationship between local and metacommunity stability as influenced by dispersal. The baseline dispersal rate *d* and the body size scaling coefficient *z* determine the species-specific dispersal rate 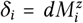, where *M_i_* is the mass of species *i*. All species have the same dispersal rate *d* when *z* = 0. When *z* is positive, larger bodied species have faster dispersal rates, whereas when *z* is negative, it is smaller bodied species that have faster dispersal rates. Other parameters vary across all simulations as indicated by Table 1. Error bars denote ± 2 S.E.M., and are too small to see.

Metacommunities are most likely to be stable when larger-bodied species disperse through spatial networks of habitat patches faster than smaller-bodied ones (that is, for *z* > 0, Fig. 2). Thus, having more patches with conditions that promote stability and experiencing higher overall spatial coupling both stabilize metacommunities. For a given proportion of stable local food webs and for all but the strongest global coupling (*i.e.* large *d*), metacommunities with a positive relationship between body size and dispersal rate are most likely to stable, while those with negative scaling are least likely to be stable.

The stabilizing effects of positive body size-dispersal scaling are most pronounced when the global link strength among habitat patches *d* is relatively low. When *d* =1, metacommunity stability with size-dependent dispersal is nearly indistinguishable from the case of uniform dispersal. As overall large levels of dispersal provide a substantial stabilizing effect, positive body size-dispersal scaling therefore appears to be beneficial for stability because large consumers increase spatial connectivity above the baseline rate set by primary producers (*i.e.* it is guaranteed that *δ_i_* ≥ *d*). With negative scaling, these same consumers have lower dispersal rates than producers (i.e. *δ_i_* ≤ *d*), lowering the overall spatial coupling between metacommunity patches and reducing stability.

Correlations between key model parameters (see Methods) and metacommunity stability are shown in Fig. 3, confirming the importance of dispersal rules. Of all model parameters examined, the baseline dispersal rate *d* and the body size scaling parameter *z* have by far the strongest stabilizing effects. In contrast, species richness *S* and food web connectance *C* have the greatest de-stabilizing effects, largely through their well-documented effects on local food webs [2, 3, 6]. No other parameter showed a notable correlation with stability, either positive or negative.

**Figure 3:**
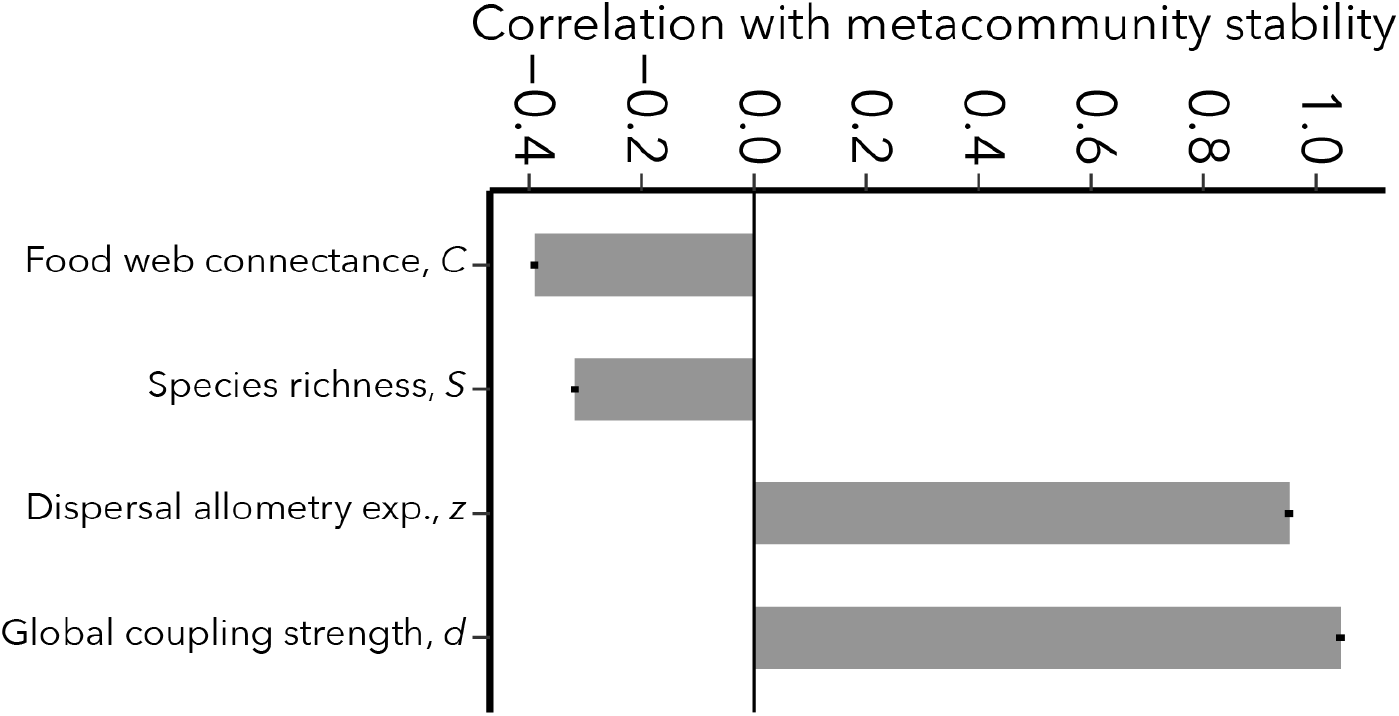
The effects of key parameters on metacommunity stability. Stability is as defined in Fig. 2 and parameters are defined in Table 1. Correlations given are coefficients from the best-fitting generalized linear model ±2 SEM. Positive correlations indicate that larger values of a parameter correspond to a higher probability that a randomly-assembled metacommunity will be stable.

The results in Fig. 2 show how body size scaling of dispersal can stabilize metacommunities by increasing coupling among patches. In Fig. 4, we compare stability of metacommunities with body size-dispersal scaling to those where the same set of dispersal rates are randomly re-assigned to species (permuted metacommunities). With these comparisons we ask whether, for a given global link strength *d*, it is coupling by large-bodied species specifically or simply greater coupling overall that drives stability. The 1:1 line in Fig. 4 shows where the leading eigenvalue *λ*_1_ of a metacommunity is the same as median leading eigenvalues of the corresponding permuted metacommunities. Most eigenvalue comparisons fall near this line across the entire range of dispersal scaling exponents *z* and global link strengths *d* (Supplementary Fig. 1). Figure 5 presents these results categorized by their qualitative effects on stability. The sign of the leading eigenvalues for the metacommunities with permuted dispersal are typically the same as the intact ones, indicating that the median effect of shuffling dispersal rates among species is rarely a qualitative change in stability. In fact, there is no combination of body size scaling *z* and dispersal coupling *d* where a qualitative change in stability occurs in more than 25% of metacommunities (Supplementary Fig. 2). Yet despite the frequent preservation of qualitative dynamics, re-arranging which species exhibit the highest dispersal rates on a patch network does show a high potential to alter the magnitude of *λ*_1_ (Fig. 4; Supplementary Figs. 1 & 2) and therefore the rates at which metacommunities return to, or depart from, steady states following perturbation. We conjecture that dispersal hierarchies may be important for the transient dynamics of ecosystems, and suggest that an understanding of how dispersal rules impact different dimensions of ecological stability will be an important goal for future work.

**Figure 4:**
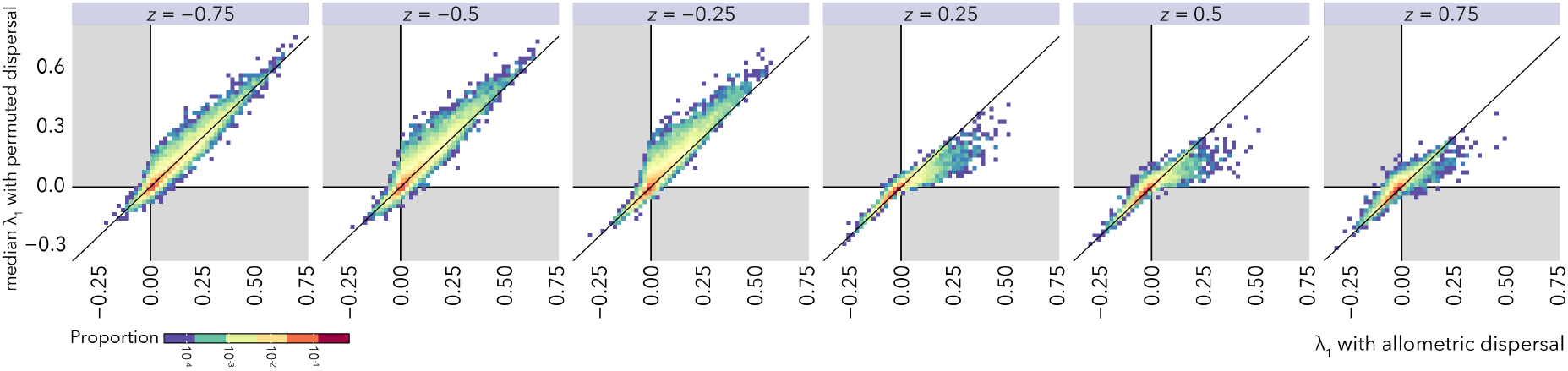
The effect of dispersal variation on metacommunity stability for spatial link strength *d* = 0.1. The metacommunity is stable when the real part of the leading eigenvalue of the metacommunity Jacobian *λ*_1_ < 0. Allometric dispersal is defined eq. 2. Permuted dispersal refers to cases where allometric dispersal rates were randomly reassigned to new species. Each unique metacommunity with allometric dispersal was compared to 100 randomly permuted counterparts, and are shown with 1:1 lines. Grey regions mark portions of the plot representing qualitative changes in stability.

**Figure 5:**
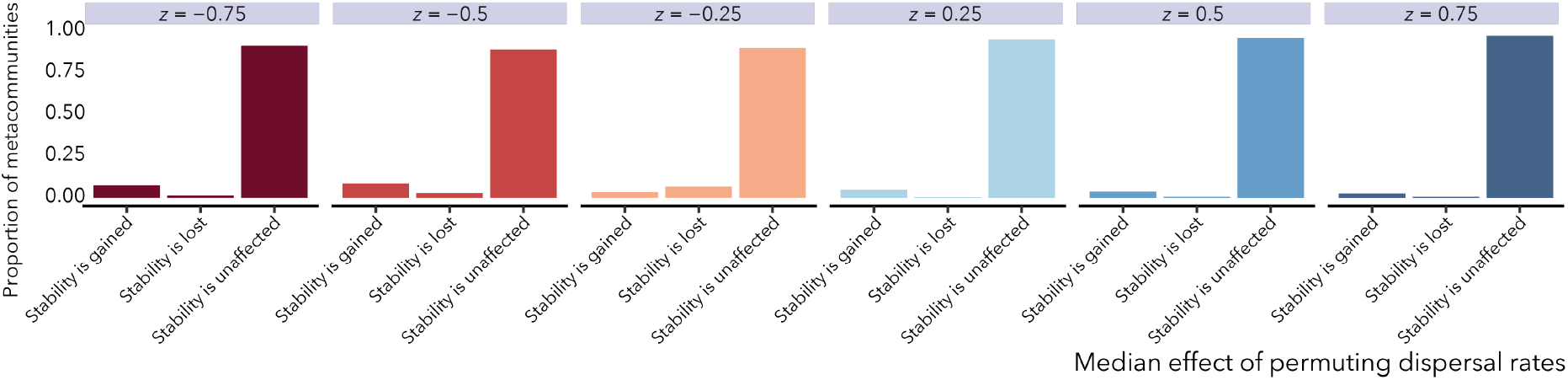
Eigenvalues *λ*_1_ from fig. 4 categorized by qualitative effects on stability. Categories *Stability is gained* and *Stability is lost* correspond to cases where the median effect of permuting dispersal rates is a change in the sign of *λ*_1_. *Stability is unaffected* indicates no sign change.

## Discussion

Our model reveals an apparent stabilizing effect of positive body size scaling of dispersal in heterogeneous metacommunities (Fig. 2), consistent with some conceptual and mathematical theories [14, 16, 61]. However, by comparing metacommunities with allometric dispersal to those whose dispersal rates were random permutations of the original values (Figs. 4 & 5), we found evidence that increased overall connectivity, and not predator movement *per se*, was responsible for metacommunity stability. Increased connectivity manifests either through larger values of global patch coupling *d* or stronger positive body size-dispersal scaling *z*, and generates conditions in which fewer stable patches are required to “rescue” unstable patches in a metacommunity.

There are many potential explanations for why random dispersal maintains roughly the same degree of metacommunity stability as positive body size scaling. The stabilizing effects of dispersal in randomly assembled metacommunities have been shown to operate when dispersal is both homogeneous and variable among species [36]. We conjecture that, for the non-random food webs we examine here, the existence of stabilizing structural motifs could be enhanced by dispersal. Consumers play key roles in the stabilizing effects of food webs compartments [62], including linking separate energy channels [16, 63] and forming long interaction loops [64]. Higher dispersal by these consumers, even if they are not top predators, could substantially enhance metacommunity stability by connecting stabilizing subscommunities across habitats. Additionally, patterns of dispersal where low level consumers have the greatest dispersal connectivity have been shown to confer stability in some tri-trophic metacommunities [54]. Finally, differential dispersal among species at similar trophic levels could generate competition-colonization and fecundity-dispersal trade-offs (reviewed in [65]), which may enhance stability in simple metacommunities. How the above mechanisms operate in more complex spatial food webs remains an open question.

Negative body size-dispersal scaling relationships lead to lower stability in our model by lowering dispersal coupling relative to situations where all species in the food web share the same dispersal rate. These results are most likely relevant for systems where connectivity results from passive transport of smaller-bodied organisms, for example wind-dispersed plants or freshwater zooplankton, flooding-dispersed pond invertebrates, and marine organisms with planktonic larvae. Passive dispersal can generate stochastic variation in connectivity [66], which may have complicated effects on stability. Yet it can also lead to increased connectivity by large numbers of individuals [49, 67, 68]. Thus, while negative scaling could potentially equate to low stability, the overall levels of dispersal may be very high in systems where lower trophic levels disperse at greater rates, reducing the scaling effect and leading to higher stability.

Spatial variation in model parameters were central to how dispersal connectivity influenced stability in our model. The positive relationship between stability and the dispersal parameters *d* and *z* was consistent throughout the parameter region we explored; additional numerical results not shown confirmed this relationship more broadly. This pattern contrasts with some other metacommunity models that exhibited a unimodal relationship between dispersal and stability [69, 70, 71, 72]. Uni-modal dispersal-stability relationships occur when very high dispersal leads the system to behave as a single, well-mixed habitat. Species persistence and the variation-reducing effects of dispersal therefore no longer operate. Similar outcomes do not seem to occur in our model, at least at the parameter values examined, because rates of primary production and trophic interactions are forced to vary in space. A key effect of dispersal in the presence of such spatial variation is to reduce variation among interaction rates of the average, well-mixed food web, which actually increases stability [36, 37].

Homogenous systems could exhibit very different dynamics. Dispersal-induced instabilities are frequent in homogeneous space trophic models (e.g. [73, 13]), leading to spatially patterned steady-states or asynchronous oscillations even when local dynamics are stable. The presence of high-dispersing top predators in more complex food webs appears to generally stabilize spatially homogeneous systems [14], although faster consumer movement is also a necessary condition for spatially pattered steady states to arise in two- and three-species food chains [74, 73]. Dispersal-induced oscillatory instabilities in contrast are more likely when primary producers disperse much faster than primary and secondary consumers in simple food chains [73, 54]. Therefore, we might expect a generally stabilizing effect of positive body size scaling and a generally destabilizing effect from negative scaling with oscillatory instabilities. However, these predictions could be complicated in more complex ecological networks, where spatial dynamics can be additionally influenced by consumers sharing a common prey (e.g. [75]) or prey being consumed by a shared predator (e.g. [73]). Furthermore, pattern formation and asynchronous oscillations could impart “stability” in a different sense by lowering population variation at larger scales [54, 76].

Our definition of stability pertains to metacommunity behavior near equilibrium and is therefore limited in describing non-equilibrium dynamics. In this class of models, instabilities can indeed include trajectories that tend to zero (i.e., species extinctions), but other outcomes including non-equilibrium co-existence with synchronous or asynchronous oscillations are also possible [77, 78, 79, 80]. The relationship between species persistence and other non-equilibrium dynamics is not always clear, particularly when species interactions are nonlinear. In metacommunities, regional persistence can occur even in the presence of local extinctions [69, 70, 72]. Unstable oscillations or even local extinctions may in fact drive spatially asynchronous dynamics that enhances regional persistence [70, 54, 76, 72]. In these cases, positive body size scaling could counteract such effects; just as coupling by large, mobile consumers stabilizes heterogeneous metacommunities in our model, predator movement could dampen the amplitude of asynchronous oscillations. Wide-ranging predators may also synchronize prey dynamics [81, 82, 83]. The rich non-equilibrium behavior possible in highly speciose food webs with nonlinearities, like we examine here, will likely require extensive investigation and generalizations may be challenging [15].

Consumers, especially large-bodied top predators, are being disproportionately lost from the world’s ecosystems [61, 84, 85]. These losses have far-ranging implications for ecosystem structure and stability [86, 87, 6, 61, 84, 14]. A potential consequence of consumer extinction is loss of spatial coupling, which previous research and our results here suggest could lead to further regional instability [63, 12, 14], including increased variability and subsequent species losses. While positive body size scaling had the greatest stabilizing effect in our model, we also found that similarly strong dispersal coupling by species in other trophic positions yielded equivalent stability. Thus, our results suggest that the conservation value of connectivity may not be lost when top consumers are. Instead, identifying alternative agents of connectivity and promoting their dispersal following top consumer loss may serve as a productive strategy. In practice, finding substitutes for large bodied consumers in some ecosystems may be difficult given the out-sized role in ecosystem coupling these species play. Yet, continued threats to top consumer persistence and the potential conservation value of dispersal among other species suggest the utility of planning for robust connectivity at the community rather than population level [88, 89].

## Methods

Steady states and stability of equation (1) were studied using the generalized modeling method [6, 55, 57]. The method assumes all populations in the food web possess a steady state, allowing us to re-cast population densities and functions as normalized proportions of the steady state. Thus, for each population 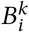 of *i* = 1,…, *S* species across *k* =1,…, *N* patches there exists a steady state 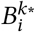 that allows us to define the normalized densities 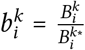. The normalized equations for the non-dispersal components of the metacommunity are therefore

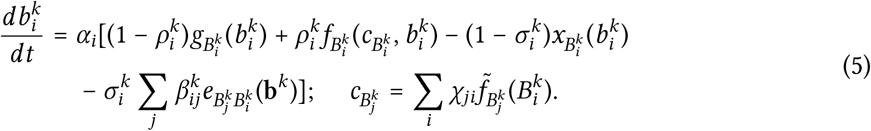

The functions *g, f, x*, and *e* represent the functions *G, F, X*, and *E* from equation (1) normalized by their values evaluated at steady state; the newly introduced variables 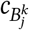 and 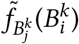 represent the total amount of food available to species *j* and the contribution of species *i* to the food available to species *j*, respectively [56]. Additional scale and branching parameters that arise from the normalization procedure set the rates and strength of interactions in the local food web. The normalized turnover rate *α_i_* scales the biomass flow rates for each species in the food web with mass, 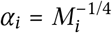, while the branching parameters quantify the structure of these flows. Specific interpretations of the scale and branching parameters are given in Table 1 and provided in detail in [55].

Determining stability of equation (4) relies on establishing the Jacobian matrix for the local food web, **J**_*k*_. The Jacobian is constructed from elements that describe the change in the dynamic equation for each species that occurs given a change in each component state variable and the functions of each state variable near steady state. The local food web Jacobian therefore is a square *S* × *S* matrix where diagonal entries *J_i,i_* describe the effects of a change in species *i* on itself and the non-diagonal entries *J_j,i_* describe the effects of species *i* on species *j*. Quantifying the changes of functions of state variables near steady state is accomplished in the generalized modeling framework by defining the following exponent parameters [6, 13, 55, 56, 57],

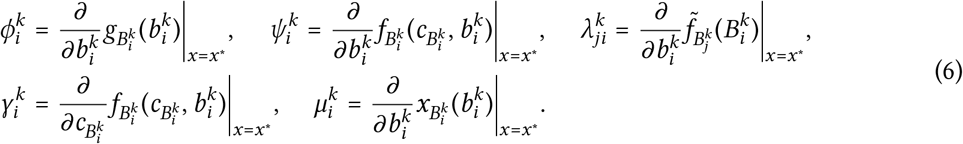

These exponent parameters can be interpreted formally as elasticities [56, 57]. Furthermore, they recapitulate effects of relevant non-linearities on ecological dynamics commonly employed in standard ecological models such as the amount of saturation in the functional response 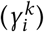, the shape of the producer growth function 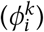, the presence of intraspecific consumer interference 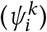, and the density-dependence of consumer mortality rates 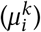. The exponent parameter 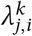 can be interpreted as the adaptability of consumer preferences to different prey items; following previous work [6] we assume constant preferences for simplicity.

Niche model food web topologies that specify the structure of interactions in equation (5) were generated following [18]. Each species *i* is assumed to exist on a niche axis between [0,1] and is assigned a niche value using a uniform distribution. The species then is assumed to consume all species over a range *r_i_* that is near or below the position of species *i* on the niche axis, generating a trophic hierarchy. The location of the range is assigned using a beta function with expected value 2*C*. We chose input values for species richness *S* as integers between 10 and 30 inclusive, [10..30], and for connectance *C* as (0.12,0.14,…, 0.24). The latter range was chosen to encompass empirically observed connectance values [90]. Only webs with a single connected component were retained for analysis.

Spatial networks were generated as random geometric graphs (RGG). Networks of *N* patches were generated by first randomly assigning coordinates in 2 dimensional space to each patch drawn from a standard uniform distribution. patches were then connected if the Euclidean distance between their coordinates fell below a threshold *n*. Networks with unconnected patches were discarded. For results included here, we used *N* = 10 and *n* = 0.32, yet simulations with greater numbers of patches, different values of *n*, and even different network configurations exhibited qualitatively similar results.

The spatial structure of the RGG is encoded in the Laplacian matrix **L**. The Laplacian **L** is an *N* × *N* matrix where the diagonal entries *L*_*k*=*l*_ represent the number of dispersal connections (i.e. degree) of each patch *k*. The off-diagonal elements of *L* reflect dispersal connections between individual patches, where *L*_*k*≠*l*_ = −1 when patches *k* and *l* are connected and 0 otherwise. Linking the Laplacian to the dispersal matrix **D** yields the spatial structure of the dispersal network. The dispersal matrix is an *S* × *S* matrix with species-specific dispersal rates on the diagonal *D*_*i*=*j*_ = *δ_i_* and all other entries zero. Using **L** ⊗ **D**, with ⊗ being the Kroenecker product, yields the *SN* × *SN* block matrix that describes the pattern of connections between all patches by all species.

Local food webs were embedded in the spatial structure of the metacommunity using equation (3). Each local web Jacobian **J**_*k*_ was numerically placed on the diagonal of the metacommunity food web matrix *P*. Because we assume environmental heterogeneity, the entries of each **J**_*k*_ in a given *P* vary, although the topologies remain fixed. Environmental heterogeneity is implemented as variation in branching and elasticity parameters; the value of each parameter for each species in a local web **J**_*k*_ was determined by drawing from uniform distributions in the appropriate range defined in Table 1.

We generated one hundred unique food web topologies for each combination of species richness *S* and connectance *C*, yielding 7,700 unique metacommunity topologies. We explored metacommunity dynamics over five values of dispersal coupling *d* and seven values of dispersal allometry *z*, yielding 192,500 unique metacommunities. Dispersal structure of each unique metacommunity was then permuted 100 times; the stability of the metacommunity was compared with the median stability of the random dispersal metacommunities.

Associations between different parameters and metacommunity stability were quantified using generalized linear models (GLMs) with binomial errors and a logit link function, with an information criterion-based model selection scheme. Following White et al. [91] we use GLMs as a framework for partitioning variance and correlations between important model parameters and metacommunity stability, and not for assessing statistical significance. We first fitted a global model comprising linear combinations of fixed effects for local food web species richness *S*, web connectance *C*, the strength of spatial network links *d*, the body size scaling exponent for allometric dispersal *z*, and the spatial variances and means of parameters describing predator satiation, interaction strengths, and nutrient availability to primary producers (Table 1). We then computed Akaike’s information criterion (AIC) for all submodels comprising different combinations of fixed effects in the global model, and selected the model with the lowest AIC score as the best fitting. This model included four terms: species richness *S*, web connectance *C*, global spatial network link strength *d*, and the exponent of allometric dispersal *z* (Fig. 3).

## Acknowledgments

K.E.A. was supported by a grant from the National Science Foundation #DEB-1553718. A.K.F. was supported by a fellowship from the National Research Council Research Associateship Program.

## Authors contributions

K.E.A. and A.K.F. conceptualized the study, wrote the manuscript, and contributed analyses.

