## Supplemental Figures for "Body size dependent dispersal influences stability in heterogeneous metacommunities"

### Supplementary figures

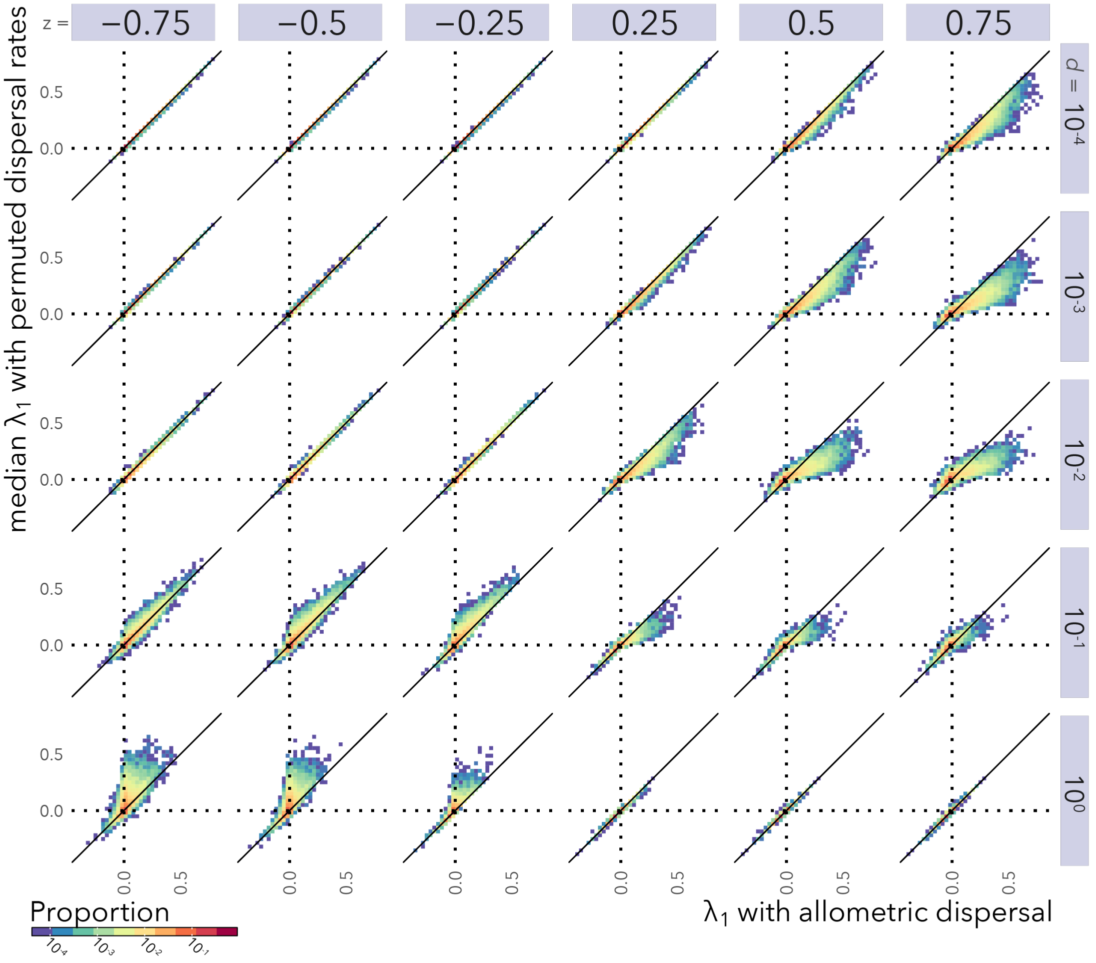


Figure S1: The effect of dispersal variation on metacommunity stability for all combinations of $d$ and $z$ (see Main Text). The metacommunity is stable when the real part of the leading eigenvalue of the metacommunity Jacobian $\lambda_1 < 0$. Allometric dispersal is defined eq. 3 of the Main Text. Permuted dispersal refers to cases where allometric dispersal rates were randomly reassigned to new species. Each unique metacommunity with allometric dispersal was compared to 100 randomly permuted counterparts, and are shown with 1:1 lines.

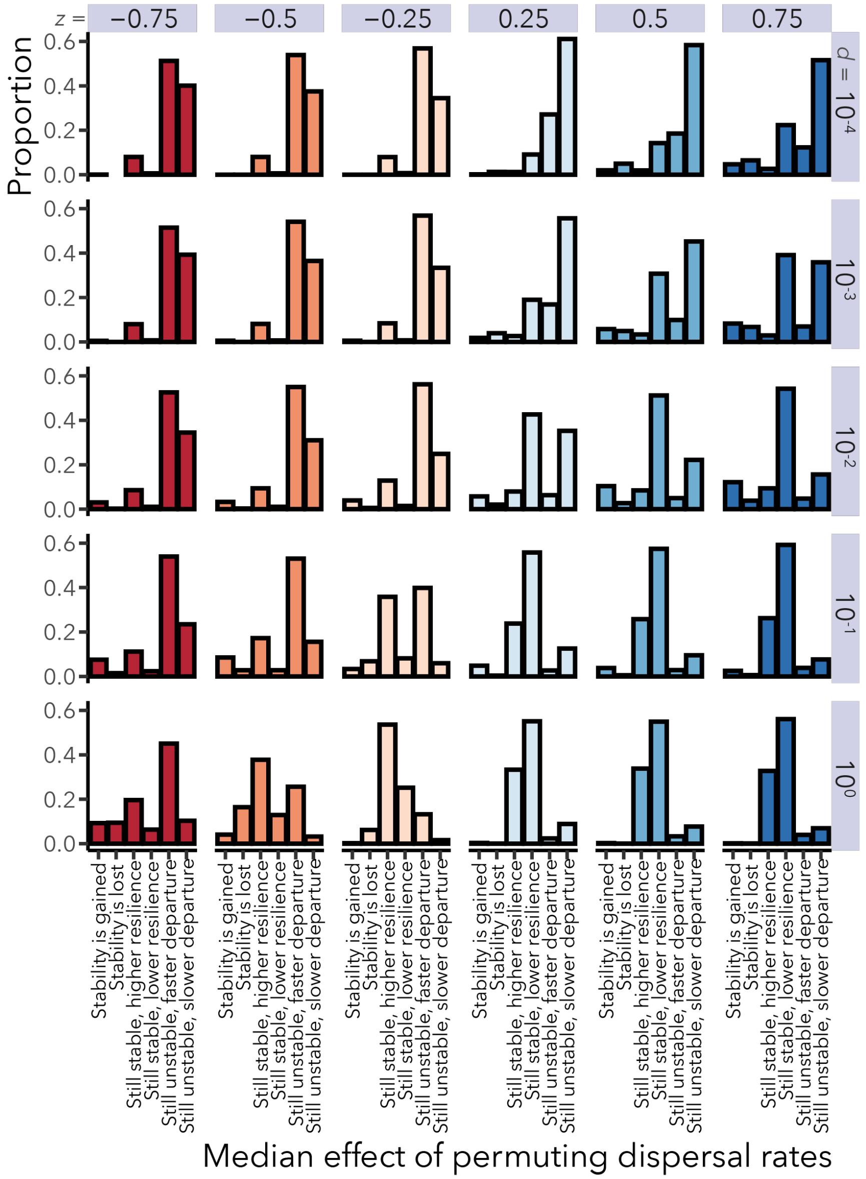


Figure S2: Eigenvalues $\lambda_1$ from Fig. S1 categorized by qualitative effects on stability for all combinations of $d$ and $z$ (see Main Text). Categories *Stability is gained* and *Stability is lost* correspond to cases where the median effect of permuting dispersal rates is a change in sign of $\lambda_1$. The remaining categories refer to cases where the magnitude, but not the sign, of the median $\lambda_1$ changes, leading to faster or slower departures from (or returns to) an unstable (or stable) steady state.
